# Sharpening of hierarchical visual feature representations of blurred images

**DOI:** 10.1101/230078

**Authors:** Mohamed Abdelhack, Yukiyasu Kamitani

**Author notes:** Correspondence should be addressed to: Yukiyasu Kamitani, Ph.D., Graduate School of Informatics, Kyoto University, Yoshida-honmachi, Sakyo-ku, Kyoto 606-8501, Japan.

## Abstract

The robustness of the visual system lies in its ability to perceive degraded images. This is achieved through interacting bottom-up, recurrent, and top-down pathways that process the visual input in concordance with stored prior information. The interaction mechanism by which they integrate visual input and prior information is still enigmatic. We present a new approach using deep neural network (DNN) representation to reveal the effects of such integration on degraded visual inputs. We transformed measured human brain activity resulting from viewing blurred images to the hierarchical representation space derived from a feedforward DNN. Transformed representations were found to veer towards the original non-blurred image and away from the blurred stimulus image. This indicated deblurring or sharpening in the neural representation, and possibly in our perception. We anticipate these results will help unravel the interplay mechanism between bottom-up, recurrent, and top-down pathways, leading to more comprehensive models of vision.

**Significance statement:** One powerful characteristic of the visual system is its ability to complement visual information for incomplete visual images. It operates by projecting information from higher visual and semantic areas of the brain into the lower and mid-level representations of the visual stimulus. We investigate the mechanism by which the human brain represents blurred visual stimuli. By decoding fMRI activity into a feedforward-only deep neural network reference space, we found that neural representations of blurred images are biased towards their corresponding deblurred images. This indicates a sharpening mechanism occurring in the visual cortex.

## Introduction

Perception is the process by which humans and other animals make sense of the environment around them. It involves integrating different sensory cues with prior knowledge to arrive at a meaningful interpretation of the surroundings. This integration is achieved by means of two neuronal pathways: a bottom-up stimulus driven pathway, which processes sensory information hierarchically, and an intrinsic pathway, which performs both recurrent processing through lateral pathways and projection of prior information down the hierarchy (we call it top-down pathway for abbreviation; Friston, 2005; Arnal and Giraud, 2012; Clark, 2013; Summerfield and de Lange, 2014; Heeger, 2017). The interplay mechanism between these two pathways is still an open question.

Previous studies have given rise to two main hypotheses to explain the top-down modulation process. The sharpening hypothesis states that top-down signals enhance the neural representation in the lower visual areas, thus improving the quality of the degraded sensory signal (Lee and Mumford, 2003; Hsieh et al., 2010; Kok et al., 2012; Gayet et al., 2017). Conversely, the prediction error hypothesis (which originates from computer science ideas; Shi et al. (2008) states that top-down signals provide expected signal information that would be redundant if represented again in lower visual areas, and therefore gets subtracted (Mumford, 1992; Rao and Ballard, 1999). This results in an error signal that is repeatedly processed to update the prediction signal until the error signal reaches zero, which corresponds with achieving a perceptual result (Murray et al., 2002; den Ouden et al., 2009, 2012; Alink et al., 2010; Meyer and Olson, 2011; Todorovic et al., 2011; Kok et al., 2012; Gordon et al., 2017). Most recently, these two hypotheses were reconciled by a third hypothesis where prediction error is computed to be later used to sharpen the neural representations (Kok et al., 2012). Models of vision usually employ this mechanism to explain the interacting neural information processing pathways (Lee and Mumford, 2003; Friston, 2005; Heeger, 2017).

Most of the top-down modulation studies utilized expectation of a previously-known visual stimulus to drive the operation of top-down pathways, and have hence focused on lower visual areas. Expectation-of-stimulus tasks facilitate comparison of a visualized stimulus and an expected stimulus at the lower visual feature level. While such studies have provided an empirical framework for the operation of top-down modulation driven by expectation in the lower visual areas, they have not revealed its overall operation in regular recognition-targeting visual tasks.

In this study, we tackle this question by investigating top-down pathway operation during a natural-image visual recognition task throughout different levels of visual processing ranging from lower visual areas (V1–3) to higher visual centers (LOC, FFA, and PPA). We drive the operation of top-down modulation by applying degradation to natural images by blurring them. When visual images are degraded, the visual sensory signal is less reliable, and the visual cortex therefore depends more heavily on prior knowledge driving the top-down pathway operation. To unmask the top-down effect, we investigate how the neural representations of viewing blurred images deviate from a pure feedforward representation leading to a sharpened representation along the visual processing pathway.

To demonstrate such sharpening, we measured and analyzed brain activity from functional magnetic resonance imaging (fMRI) brain data from different regions of the lower and higher visual areas, to visualize the degradation effect on different levels of neural processing. We utilized deep neural network (DNN) feature space as a proxy for hierarchical representation. We used a feature decoding method devised by Horikawa and Kamitani (2017a) to map brain activity into a DNN representation space. The decoded features were analyzed for their similarity to the feedforward-only DNN features of the stimulus images and original non-blurred images. These similarities are then compared to their counterpart noisy DNN features, which account for decoding errors as a baseline for pure-feedforward behavior, to find whether predicted features deviate from the pure feedforward ones and how supplementing with prior knowledge about stimulus categories would affect the sharpening behavior. We also compared the case where the image content is successfully recognized with the one where it is not. If image sharpening were in operation, it would be expected that the top-down effect would be boosted due to successful perception.

## Methods and materials

### Subjects

Five healthy subjects (three males and two females, aged between 22 and 33) with normal or corrected-to-normal vision took part in the fMRI experiments. The study protocol was approved by the Ethics Committee of ATR. All the subjects provided written informed consent for their participation in the experiments.

### Visual stimuli

Both original and blurred image stimuli were shown. The images were selected from the ImageNet online database (Jia Deng et al., 2009), which is the database used for training, testing, and validation of the pre-trained DNN model used in this study (see below). The database contains images that are categorized by a semantic word hierarchy organized in WordNet (Fellbaum, 2012). First, images with a resolution lower than 500 pixels were excluded, then the remaining images were further filtered to select only those that showed the main object at or close to the midpoint of the image. The selected images were then cropped to a square that is centered on the midpoint. If no acceptable image remained after this filtration process, another image was obtained from the worldwide web through an image search.

We created three different levels of blurring for the blurred image stimuli. Blurring was conducted by running a square-shaped averaging filter over the whole image. The size of the filter relative to the image size dictated the degradation level. The three degradation filters used had a side length of 6%, 12%, and 25% of the side length of the stimulus image. We then added the original stimulus image represented by a level of 0% (Figure 1A).

**Figure 1.**
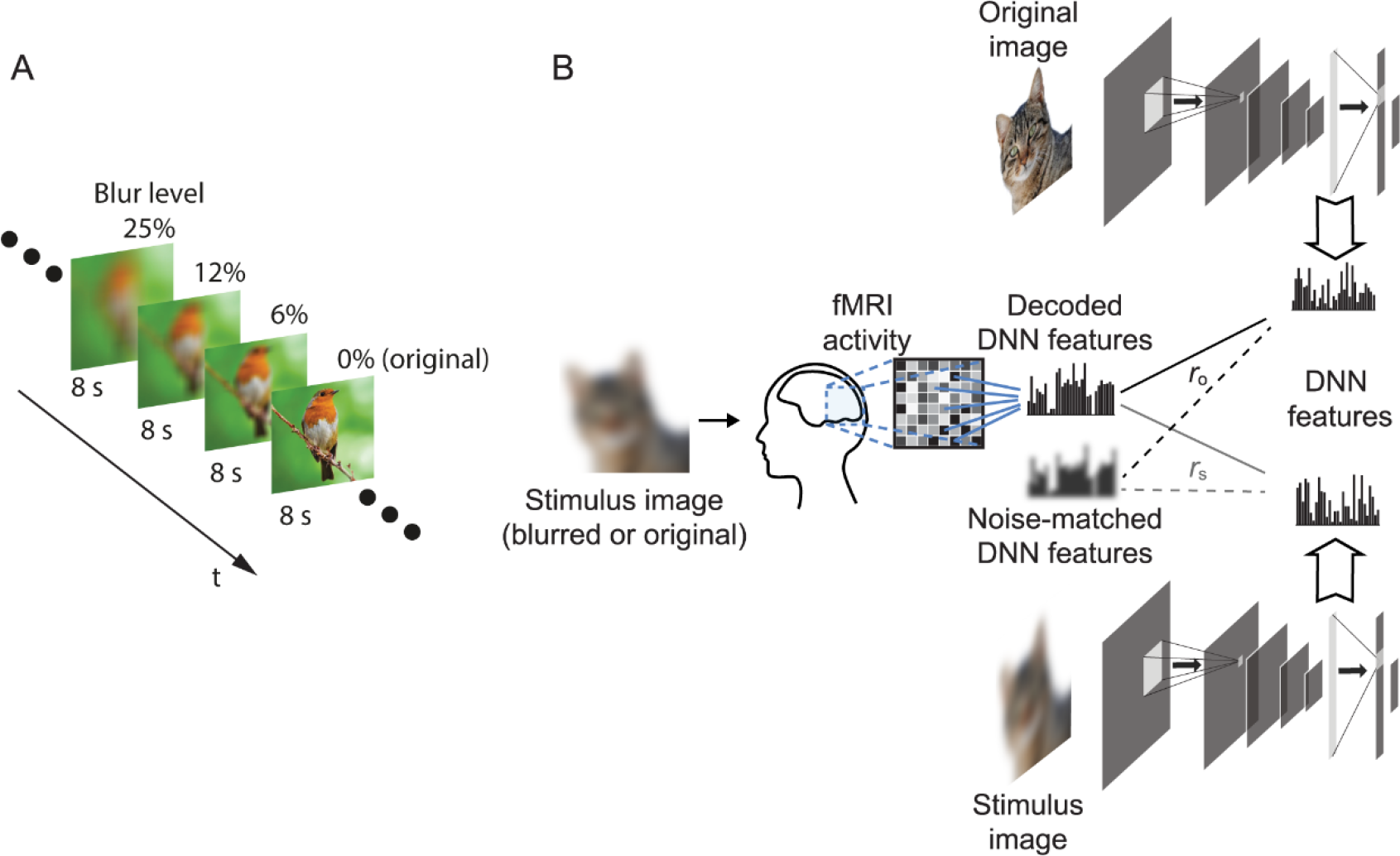
Study design. (A) The stimulus sequence was divided into sequences of four stimuli each. Stimuli in the same sequence contained different blur levels of the same image organized from the highest blur level (25%) to the lowest (0%). Each stimulus was presented for 8 seconds. (B) Overview of the feature decoding analysis protocol; fMRI activity was measured as the subjects viewed the stimulus images presented, described in A. Trained decoders were used to predict DNN features from fMRI activity patterns. The decoded features were then analyzed for their similarity with the true DNN features of both the original image (*r*_o_) and stimulus image (*r*_s_). The same procedure was also conducted for noise matched DNN features that are composed of true DNN features with additional Gaussian noise to match predicted features from fMRI.

### Experimental design

Experiments were divided into: 1) the decoder training runs where natural undegraded images were presented, and 2) the test image runs where the blurred images were presented. Images included in the training and test datasets were mutually exclusive. In the decoder training runs, we selected one stimulus image for each of the AlexNet classification categories defined in the last layer, resulting in a total of 1000 stimulus images. This training stimulus set selection was conducted to avoid any bias to certain categories in the decoder. This dataset was divided into 20 runs of 50 images each. The subject was instructed to press a button when the image was a repeat of the image shown one-back. In each run, 5 of the 50 images were repeated in the following trial to form the one-back task. Each image was shown once to the subject (except for the one-back repetitions).

The test image runs consisted of two conditions. In the first condition, the subjects did not have any prior information about the stimuli presented (no-prior condition). In the second condition, the subjects were provided a semantic prior in the form of category choices (category-prior condition). The stimuli in the category-prior condition consisted of images from one of five object categories (airplane, bird, car, cat, or dog). The subject was informed of these categories prior to the experiment, but not the order in which they were to be presented.

The stimuli in both of the test conditions were presented in sequences of maximum blurring to original image (25%, 12%, 6%, and 0% blurring). Each sequence consisted of stimuli representing all four levels of blurring of the same original image. We selected this order of presentation to avoid the subjects having a memory-prior of the less blurred stimuli when viewing the more blurred ones. For each condition, the sequences for 80 images were randomly distributed across two runs (40 images each). The runs belonging to the same test experimental condition were conducted in the same experimental session. The training and test experiments were conducted over the course of five months in total for all subjects.

All image presentation was performed using Psychtoolbox (Kleiner et al., 2007). Each image (12 × 12 degrees) was presented in a flashing sequence for 8 s at 1 Hz (500 ms on time). Images were displayed in the center of the display with a white central fixation point. The fixation point changed from white to red 500 ms before each new stimulus appeared. A 32-s pre-rest and 6-s post rest period were added at the beginning and end of each run respectively. Subjects were required to fixate on the central point. For test runs, subjects were required to provide voice feedback of their best guess of the perceived content of the stimulus. They were also required to report the certainty level of that guess by pressing one of two buttons, one indicating certainty and the other indicating uncertainty. We checked if the vocal reports caused excessive motion by the subject that leads to degradation in the data quality but found that the motion correction results were comparable to runs without vocal response by the same subjects.

### MRI acquisition

FMRI data was collected using a 3-Tesla MAGNETOM Verio (Siemens Medical Systems, Erlangen, Germany) MRI scanner located in Kokoro Research Center, Kyoto University. For image presentation experiments, an interleaved T2^*^-weighted multiband accelerated EPI scan was performed to obtain images covering the whole brain. The scanning parameters were TR = 2000 ms; TE = 43 ms; flip angle = 80°; FOV = 192 × 192 mm; voxel size = 2 × 2 × 2 mm; slice gap = 0 mm; number of slices = 76; multiband factor = 4. For localizer experiments, an interleaved T2*-weighted gradient-EPI scan was performed with the following parameters TR = 3000 ms; TE = 30 ms; flip angle = 80°; FOV = 192 × 192 mm; voxel size = 3 × 3 × 3 mm; slice gap = 0 mm; number of slices = 46. For retinotopy experiments, an interleaved T2*-weighted gradient-EPI scan was also performed where the scanning parameters were TR = 2000 ms; TE = 30 ms; flip angle = 80°; FOV = 192 × 192 mm; voxel size = 3 × 3 × 3 mm; slice gap = 0 mm; number of slices = 30. T2-weighted turbo spin echo (TSE) images with the same slice positions as the EPI images were also acquired, to act as high-resolution anatomical images. The parameters for the anatomical sequences matching the image presentation acquisition were TR = 11 000 ms; TE = 59 ms; flip angle = 160°; FOV = 192 × 192 mm; voxel size = 0.75 × 0.75 × 2.0 mm; slice gap = 0 mm; number of slices = 76. For the localizer experiment, the TSE parameters were TR = 7020 ms; TE = 69 ms flip angle = 160°; FOV = 192 × 192 mm; voxel size = 0.75 × 0.75 × 3.0 mm; slice gap = 0 mm; number of slices = 48. For the retinotopy TSE acquisition the parameters were TR = 6000 ms; TE = 58 ms; flip angle = 160°; FOV = 192 × 192 mm; voxel size = 0.75 × 0.75 × 3.0 mm. T1-weighted magnetization-prepared rapid acquisition gradient-echo (MP-Rage) fine-structural images of the entire head were also acquired. The scanning parameters for these were TR = 2250 ms; TE = 3.06 ms; TI = 900 ms; flip angle = 9°; FOV = 256 × 256 mm; voxel size = 1 × 1 × 1 mm number of slices = 208.

### MRI data preprocessing

After rejection of the first 8 seconds of each acquisition to avoid scanner instability effects, the fMRI scans were preprocessed using SPM8 (http://www.fil.ion.ucl.ac.uk/spm, RRID: SCR_007037), including 3D motion correction, slice-timing correction, and co-registration to the appropriate high resolution anatomical images. Both scans were then also co-registered to the T1 anatomical image. The EPI data were then interpolated to 2 × 2 × 2 mm voxels and further processed using Brain Decoder Toolbox 2 (https://github.com/KamitaniLab/BrainDecoderToolbox2, RRID: SCR_013150). Volumes were shifted by 2 s (1 volume) to compensate for hemodynamic delays, then the linear trend was removed from each run and the data were normalized. As each image was presented for 8 s, it was represented by four fMRI volumes. These four volumes were then averaged to provide a single image with increased signal to noise ratio for each stimulus image. The averaged voxel values for each stimulus block were used as an input feature vector for the decoding analysis.

### Region of interest construction

Regions of interest (ROIs) were created for several regions in the visual cortex, including the lower visual areas V1, V2, and V3, the intermediate area V4, and the higher visual areas consisting of the lateral occipital complex (LOC), parahippocampal place area (PPA), and fusiform face area (FFA).

First, anatomical 3D volumes and surfaces were reconstructed from T1 images using the FreeSurfer reconstruction and segmentation tool (https://surfer.nmr.mgh.harvard.edu/, RRID: SCR_001847). To delineate the areas V1–4, a retinotopy experiment was conducted following a standard protocol (Engel et al., 1994; Sereno et al., 1995) involving a rotating double wedge flickering checkerboard pattern. The brain activity data for this experiment was analyzed using the FreeSurfer Fsfast retinotopy analysis tool (https://surfer.nmr.mgh.harvard.edu/fswiki/FsFast, RRID: SCR_001847). The analysis results were visually examined and ROIs were delineated on a 3D inflated image of the cortical surface. Voxels comprising areas V1–3 were selected to form the lower visual cortex (LVC) ROI.

Functional localizer experiments were conducted for the higher visual areas. Each subject undertook eight runs of 12 stimulus blocks. For each block, intact and pixel-scrambled images of face, object, and scene categories were presented in the center of the screen (10 × 10 degrees). Each block contained 20 images from one of the previous six categories. Each image was presented for 0.3 s followed by 0.45 s of blank gray background. This led to each block having a duration of 15 seconds. Two blocks of intact and scrambled images of the same category were always displayed consecutively (the order of scrambled and intact images was randomly chosen), followed by a 15- s rest period with a uniform gray background. Pre-rest and post-rest periods of 24 s and 6 s respectively were added to each run. The brain response to the localizer experiment was analyzed using the FreeSurfer Fsfast event related analysis tool. Voxels showing the highest activation response to intact images for each of the face, scene, and object categories in comparison with their scrambled counterparts were visualized on a 3D inflated image of the cortical surface and delineated to form FFA, PPA, and LOC regions respectively. Voxels constituting the areas FFA, PPA, and LOC were then selected to form the higher visual cortex (HVC) ROI and the aggregation of LVC, V4, and HVC was used to form the visual cortex (VC) ROI. Selected ROIs for both retinotopy and localizer experiments were transformed back into the original coordinates of the EPI images.

### Deep neural network model

The neural representations were transformed into a deep neural network (DNN) feature proxy using the AlexNet DNN model (Krizhevsky et al., 2017). The Caffe implementation of the network packaged for the MatConvNet tool for MATLAB (Vedaldi and Lenc, 2015) was used for implementation. This network was trained to classify 1000 different image categories with images from the ImageNet database. The model consisted of 8 layers; the first five of which were convolutional layers, while the last three were fully-connected layers. The input to each layer is the output of the previous one as follows

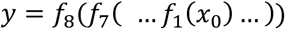

where x is the input image and y is the resulting image classification vector and the function *f*_*n*_ is the operation for each layer is

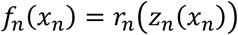

and

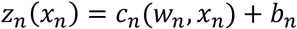

where *r*_*n*_ is a non-linearity function of the n^th^ layer (rectified linear operation for the first seven layers and softmax for the final layer), *w*_*n*_ is the n^th^ layer weight matrix that are pretrained in the model using the ImageNet dataset, *c*_*n*_ is the operation conducted at the n^th^ layer between its input and weights (convolution in the case of convolutional layers and matrix multiplication in the case of fully-connected layers), and *b*_*n*_ is the n^th^ layer bias term. We extract features from each layer *n* as the output of *z*_*n*_(*x*_*n*_) before the application of the non-linearity.

One thousand features were extracted from each layer (out of 290400, 186624, 64896, 64896, 43264, 4096, 4096, and 1000 features from DNN layer 1–8 respectively), with the features with the highest feature decoding accuracy according to the mean accuracy of the five subjects’ data in Horikawa and Kamitani (2017a) being selected. All the feature units in the last layer were selected, as this layer contained 1000 units in total. The features from each layer were labelled as DNN1–DNN8.

### DNN feature decoding

Multiple linear regression decoders were constructed to predict each feature extracted from the voxels of each ROI from the layers of the DNN. The decoders were constructed using sparse linear regression (SLR; Bishop, 2006). This algorithm assumes that each feature can be predicted using a sparse number of voxels and selects the most significant voxels for predicting the features (For details, see Horikawa and Kamitani, 2017a).

A decoder was constructed for each feature. Voxel selection was undertaken for each ROI, to select the 500 voxels with the highest correlations with each feature value. FMRI data and features of the training image dataset were first normalized to a zero mean with one standard deviation. The mean and standard deviation values subtracted were also recorded. The decoders were then trained on the normalized fMRI data and DNN features. The recorded mean and standard deviation from the training fMRI data were then used to normalize the test data before decoding the features. The resulting features were denormalized only by multiplying it by the standard deviation but not the addition of the mean to avoid the effect of baseline correlation in the subsequent data analysis. For the correlation analysis, the feature vectors emanating from the DNN were normalized by subtracting the mean of the training dataset, to match the predicted features. These normalized feature vectors are referred to as “true” feature vectors in this study.

The feature pattern correlation was computed for each stimulus image by aligning the predicted 1000 features from each DNN layer and computing their Pearson correlation coefficient with the corresponding true feature vector.

### Noise-matched features

The decoded features of blurred stimulus images can be assumed to comprise the result of both bottom-up and top-down processing in addition to fMRI noise, while those of the true features from the DNN only contain the result of the bottom-up processing. To isolate the effect of the top-down processing, we defined baseline features (noise-matched feature) by adding noise to the true features. We could extract the matching noise level from the decoded features of the non-blurred stimulus images assuming that they do not elicit a sharpening top-down process and hence only contain the bottom-up and fMRI noise components. Thus, we can add noise to the true DNN features elicited from non-blurred stimulus images till their behavior matches that of the decoded features of the same images. To perform this operation, Gaussian noise was added to the true features so that the correlation between the noisy and the true features equated the correlation between the decoded and the true features. This matching noise level was calculated for each ROI/DNN layer pair in each subject.

### Feature gain

The similarity of the decoded features to the original image features and that to the stimulus image features were evaluated by the correlation coefficients, *r*_o_ and *r*_s_, respectively, and the difference was calculated

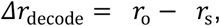

which indicates the bias toward the original features. To set a baseline, the same difference of the correlation coefficients was calculated for the noise-matched features

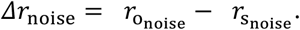

The feature gain was defined as the difference between these

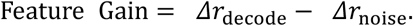

A positive feature gain means that the decoded features are more biased towards the original image features as compared to the noise-matched features.

### Content specificity

To estimate the content specificity of the predicted features from the VC, their correlation with the original image features was compared to that with the other original image features. The correlation of predicted features for each stimulus was calculated for each of the non-corresponding original images in the test dataset (n = 39), and the mean correlation was then calculated. The mean over all the stimulus images from the stimuli grouped by DNN layer with all blur levels pooled was calculated (Different image correlation), and compared with the mean of the correlations with the corresponding original images (Same image correlation). To compare this to the baseline correlation between different images, the mean of the correlation between each stimulus image vector and the feature vectors of other original images was calculated (True feature correlation).

### Behavioral data extraction

The subject vocal response was recorded manually from the voice recordings. The written record was then revised with each subject to ensure accuracy. The record was written as incorrect in the cases where the subject missed giving a voice response. In the cases when the subject missed giving a button response, the previous button response from the same sequence was used, except when the stimulus was the last (original image) or the first (most degraded image) in the sequence, when the response was set to certain and uncertain respectively. Correct responses were the ones identical to the response of the last stimulus in each sequence (original image).

### Code accessibility

The code described in the manuscript is freely available online at [https://github.com/KamitaniLab/BlurImageSharpening]. The code is available as extended data (Extended Data 1). It was created and run on MATLAB R2016b (RRID: SCR_001622) on a Linux Centos operating system on a computer cluster for parallel computing. Data to reproduce our results are also available at [http://brainliner.jp/data/brainliner/Blur_Image_Sharpening/].

## Results

### DNN feature decoding

We posed a question on how top-down modulation in the visual cortex affects the neural representation of blurred images. To address this question, we measured brain activity while presenting blurred images. The protocol involved the presentation of stimuli in blurred-to-original image sequences. Each sequence consisted of stimuli showing different blur levels of the same image presented in the order of the most blurred to the non-blurred original image (Figure 1A) so the subject is progressively receiving sharper information about the stimulus. Subjects vocally reported the perceived object in each stimulus, while also reporting their certainty of their perception. We conducted two experiments using this protocol. In the first experiment, each image (stimulus sequence) was chosen from a random object category and the subject had no prior information of the object category (no-prior condition). In the second experiment, the stimulus sequences were chosen from five predefined object categories (airplane, bird, car, cat, and dog). The subjects were informed about the object categories of the set, but not of each stimulus (category-prior condition). Using these two conditions we can analyze the effect of adding prior information on the top-down effect in different visual areas.

To examine the effect of top-down modulation, we investigated the neural representation of blurred images via the proxy of a hierarchical feedforward-only representational space (Horikawa and Kamitani, 2017a). To transform brain data into the DNN feature space, we trained multivoxel decoders to predict DNN features from brain activity data using a separate training stimulus dataset consisting of 1000 natural non-blurred images. To confirm that this choice of stimuli in training dataset did not cause the decoder to be biased to non-blurred images, we conducted a content specificity analysis as will be presented later.

Using the trained decoders, the brain activity pattern induced by each stimulus in the blurred-to-original sequences was decoded (transformed) into the DNN feature space. For each stimulus image, the Pearson correlation coefficient between its decoded feature vector and the true features of the same stimulus image (*r*_s_) at each layer was computed. In addition, the correlation between the decoded feature vector and the true features of the corresponding non-blurred original image (*r*_o_) was computed (Figure 1B). For non-blurred stimuli, *r*_s_ and *r*_o_ are identical.

### Feature gain computation

The correlation with stimulus image features (*r*_*s*_) reflects the degree to which image features resulting from feedforward processing are faithfully decoded from brain activity, while the correlation with original images (*r*_*o*_) reflects the degree to which the decoded features are “sharpened” by top-down processing, to be similar to those of the non-blurred images. Figure 2A shows a scatter plot depicting a representative result for prediction of the DNN layer 6 feature vector from the region of interest comprised of all the visual areas (Visual cortex, VC) of Subject 4. DNN6 is a higher middle layer of AlexNet where we can visualize the top-down effect on mid-level representations of visual stimuli. It is also a fully connected layer that processes global stimulus information rather than local information in the case of convolutional layers. This would lead to better separated clusters that could show the transition from the most to the least blurred. Each point represents a stimulus image pooling both category-prior and no-prior conditions, and disregarding behavioral data while the mean points are also shown (white points with black borders) to demonstrate how decreasing blur level leads to decoded features veering towards original image features. Figure 2B shows the mean of the results in Figure 2A, grouping stimuli by different blur levels. From the results of *r*_s_ and *r*_o_, we define *Δr*_decode_ as the difference between them. We notice from the representative data that decoded features have higher correlation with the original image features than with the stimulus image features, except when the blurring effect becomes too large, as in the 25% blur level. This suggests that a sharpening effect occurring in the visual cortex causes the neural representations of viewing the blurred image to mimic those of a less blurred version of it.

**Figure 2.**
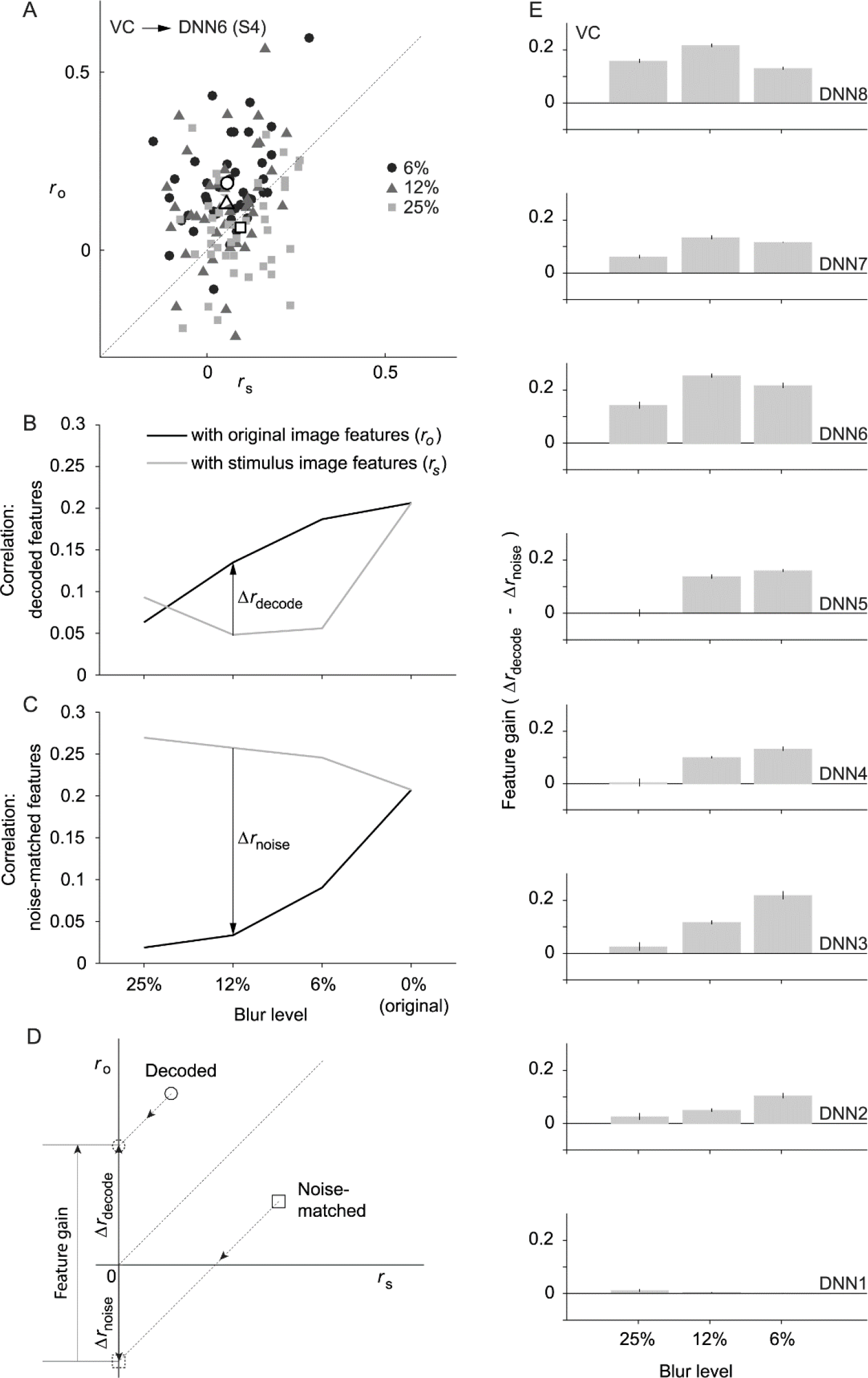
Correlation of decoded features with original and stimulus image features. (A) Scatter plot showing feature correlation of DNN6 features decoded from the whole visual cortex (VC) of subject 4, with original image features (*r*_s_; x-axis) and stimulus image features (*r*_o_; y-axis). Each point represents a stimulus image for all blurring levels except 0% while the white points with black borders show the mean of all points of the same blur level. Diagonal dotted line represents the line of equal correlation (*Δr*_decode_ = 0). (B) Representative result from DNN6 features decoded from the whole visual cortex (VC) of subject 4. Solid lines represent the mean correlation at different blur levels while pooling different experimental conditions and behavioral response data. The difference between *r*_o_ and *r*_s_ is labelled as *Δr*_decode_. (C) Representative result showing mean noise-matched feature correlation with the original and stimulus image features for different blur levels. Noise-matching was performed to match the correlation of the DNN6 predicted features of the 0% blur stimuli decoded from VC of subject 4 (thus obtaining equal values with the decoded features at the 0% level). The difference between *r*_o_ and *r*_s_ yields the noise baseline (*Δr*_noise_). (D) Feature gain is defined as the difference between *Δr*_decode_ and *Δr*_noise_. *Δr* could be defined as the displacement along the *r*_o_ axis of the point on the plot from the line of equal correlation. So by subtracting the vector representing noise matched feature correlations from decoded feature correlation, we can calculate feature gain. (E) Mean feature gain is indicated for each DNN layer for features decoded from VC at different stimulus blur levels (excluding the 0% level). Error bars indicate 95% confidence interval (CI) across five subjects.

One shortcoming of this measure (*Δr*_decode_) is that it does not have an appropriate baseline for sharpening. A value of *Δr*_decode_ equal to zero implies that decoded features are equally similar to stimulus and original image features, but it does not mean that there is no sharpening. Thus, we defined a baseline for no sharpening according to the behavior of feedforward-only processing. Decoded features from feedforward-only processing were modeled by stimulus image features plus Gaussian noise. The noise level was determined to match the decoding errors with the non-blurred images used as stimuli, in which no sharpening was assumed to be involved. Noise was added to the point where the decoded and noise-added features had nearly identical correlations to the original image features (Figure 2B and C; 0% blur level in each). The same level of noise (the mean across images in each subject and DNN layer) was added to the stimulus features of the blurred images. We then computed *r*_s_ and *r*_o_ for the noise-matched features, from which we could obtain the noise-matched baseline *Δr*_noise_ (Figure 2C).

By comparing B and C in Figure 2, it is possible to note an opposite trend in how the features are correlated. As the decoder was trained to predict image features, the natural trend for *Δr*_decode_ would be negative, similar to *Δr*_noise_. This indicates a level of alteration in the neural representation of the blurred images, to improve the match with the original images.

By subtracting the noise baseline from *Δr*_decode_, we obtained the “feature gain” incurred by top-down processing (Figure 2D). The value of the feature gain indicates how the top-down pathways affect the predicted features in comparison with pure feedforward behavior. Figure 2E shows the results of the mean feature gain for different subjects for each layer. We can observe positive significant feature gains for most of the DNN layers and blur levels (17 out of 24 DNN layer/blur level combinations; *t*-test across subjects with Bonferroni correction, *p* < 0.002, Bonferroni correction factor = 24). This suggests that top-down processing modulates neural representations to bias them towards the original images. We also noticed that the fully-connected layers DNN6–8 had more pronounced positive feature gains than the convolutional layers. Another notable issue is that the 12% blur level shows better feature gain relative to both 6% and 25% blur levels in higher visual areas. One possible explanation is that at 6% blur level the local information start to unravel leading to sharpening at the shallower layers only.

One possible cause for this result is the training scheme of the decoders that only used natural non-blurred images. This could have biased the output features to those resembling natural images. In this case, the features could be correlated to any natural image features. We investigated this possibility by measuring the content specificity of the predicted features. We computed the correlation of predicted features (excluding those with a 0% blur level) with the corresponding original image feature (*r*_o_). This was then compared with the mean correlation of the same predicted features, but with the original features of different images. This measure provided information on how tightly the predicted features were associated with the presented stimulus content, as opposed to natural images in general. Figure 3 shows the result of such a content specificity analysis. There are significant differences between correlations with the same image features and mean correlations with different image features in all layers (*t*-test across subjects, *p* < 0.05, uncorrected), indicating a tight association of the predicted features with the stimulus image content, and ruling out a decoder bias explanation.

**Figure 3.**
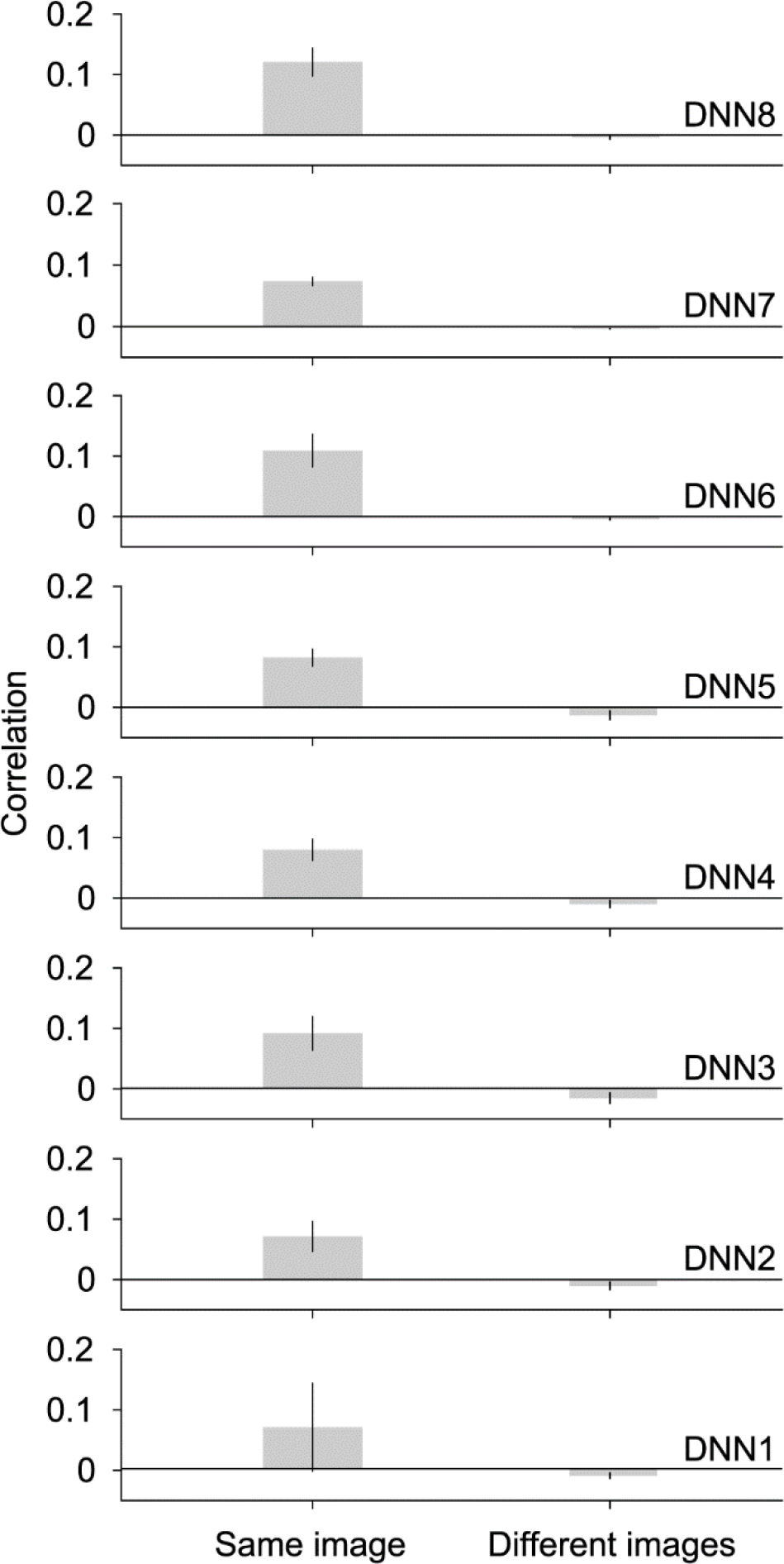
Content specificity of decoded features with blurred images. Same image correlation indicates correlation of predicted features (blur levels pooled, excluding 0%) with corresponding original image features. Different images correlation indicates the mean of correlations of the same predicted features with original image features of different images. The mean correlation is shown for different DNN layers. Error bars indicate 95% CI across five subjects.

As mentioned before, the DNN model used in this study implements hierarchical processing that is synonymous with that happening in the visual cortex. Previous studies have shown homology between the features of the DNNs and the representations in the visual cortex (Cadieu et al., 2014; Khaligh-Razavi and Kriegeskorte, 2014; Yamins et al., 2014; Güçlü and van Gerven, 2015; Horikawa and Kamitani, 2017a). To this point, we have shown the results of features predicted from the collection of all denoted visual areas (VC). We further investigated the separate visual areas of the lower, intermediate, and higher visual areas, to examine the homology between the feature gain and the visual cortex hierarchy (Figure 4). We showed that the feature gain also shows similar homologies to the visual hierarchy, in that we could observe that shallower DNN layers showed larger feature gain from the lower visual areas (V1–3), while deeper DNN layers showed larger feature gain from the higher visual areas (LOC, FFA, PPA). The results are significantly positive for most of the layers and regions of interest (ROI), especially in the higher visual areas and fully connected layers (*t*-test, *p* < 0.05, uncorrected). However, DNN1 did not show significantly positive feature gains. These results imply that feature gain also follows the same visual homology in the visual cortex areas, and that the top-down effect is more pronounced in higher visual areas.

**Figure 4.**
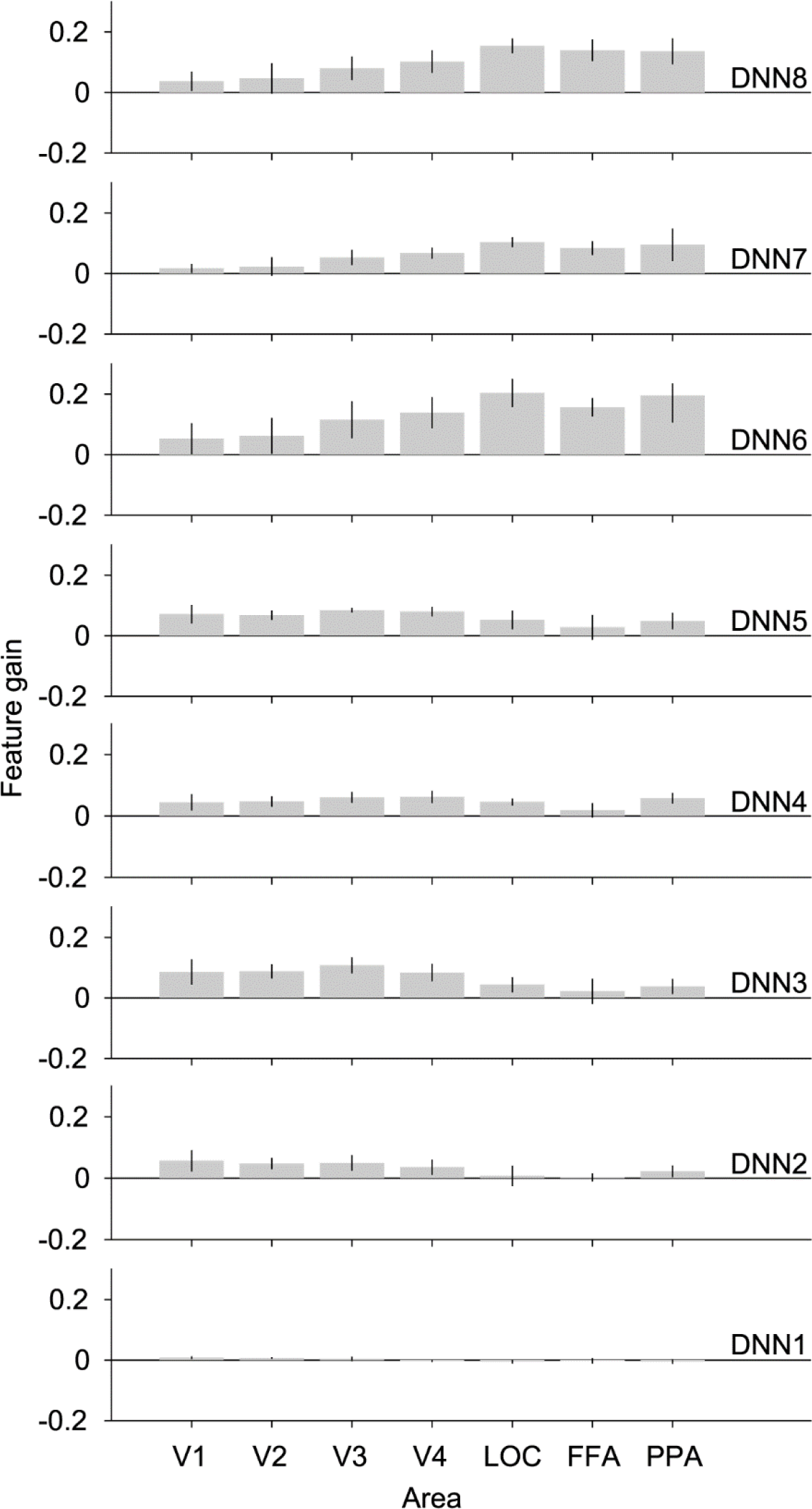
Feature gain across visual areas. Feature gain for features predicted from different visual areas. Mean feature gain is indicated for each DNN layer (blur levels pooled, 0% excluded). Error bars indicate 95% CI across five subjects.

### Effects of prior knowledge and recognition

In the previous analyses, the data from different experimental conditions were pooled together. We then further investigated the difference between the category-prior and no-prior conditions. We compared the feature gain means grouped according to the experimental condition (category-prior vs. no-prior) while pooling all the behavioral responses (Figure 5). We performed two-way ANOVA on the feature gain data using the ROI and the experimental conditions as the independent variables. The addition of a prior caused significant enhancement to the feature gain in layers DNN4, 7, 8 (*p* < 0.006, Bonferroni correction factor = 8). The difference was most pronounced in DNN8 (*p* = 0.0000026). This result indicates that addition of prior information enhances top-down modulation, thereby causing an increase in feature gain. This implies augmented sharpening of neural representations.

**Figure 5.**
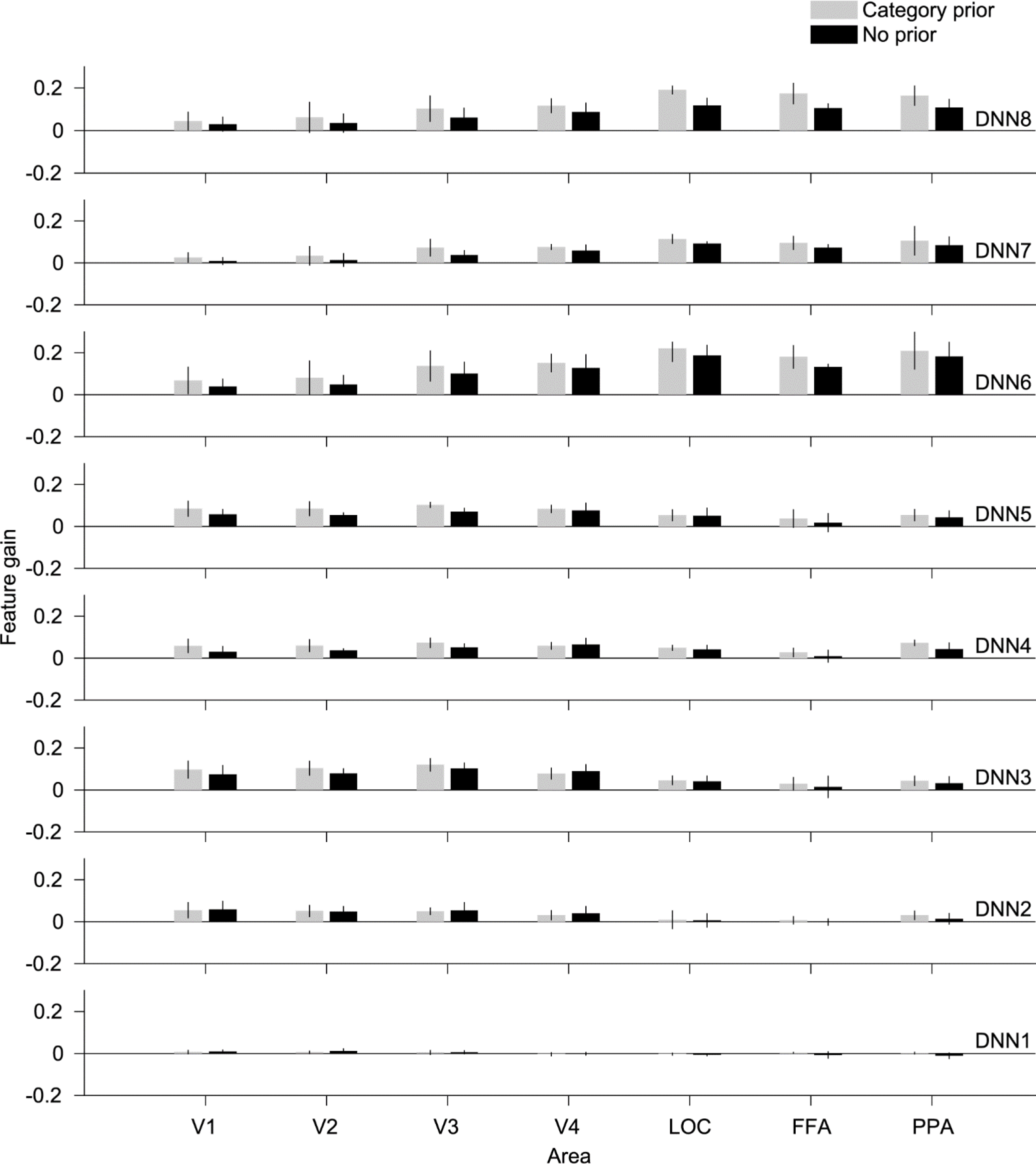
Effect of category prior. Feature gain for features predicted from different visual areas grouped by experimental condition (category-prior vs. no-prior). Mean feature gain is indicated for each DNN layer (blur levels pooled, 0% excluded). Error bars indicate 95% CI across five subjects.

This result, however, pooled both correctly and incorrectly reported results. When considering behavioral data, there are considerable differences between category-prior and no-prior conditions. The category-prior condition was characterized by a higher number of correct responses (235 out of 300 total instances for five subjects) compared with the no-prior condition (92 out of 300 total instances for five subjects). However, in the category prior condition, the task was to choose one of five categories. This could lead to false positives, as if a subject responded in a random manner, 20% of the responses would be likely to be correct. In some cases when the stimulus was highly degraded, the best guess response by the subjects could be random. To attempt to curb this problem, we could use the certainty level as an indicator of correctness, especially for the category prior. We found from the behavioral results that nearly all the trials labelled as certain were also correctly recognized (category prior: 138 out of 139 certain trials were correct; no-prior: 57 out of 70 certain trials were correct). This further supports the observation that adding priors aids recognition.

We further analyzed our data by grouping it according to both experimental condition (category-prior and no-prior) and recognition performance (correct and incorrect). We show the results of the mean feature gain over subject means for each DNN layer in Figure 6. For each experimental condition, we performed a three-way ANOVA test using ROI, recognition performance, and blur level as independent variables. For the category-prior condition, we found significant enhancement in feature gain when an image was correctly recognized in DNN6, while for the no-prior condition no significant enhancement was found (p < 0.003, Bonferroni correction factor = 16). From these results, we notice that the effect of recognition leads to a very faint enhancement in the feature gain.

**Figure 6.**
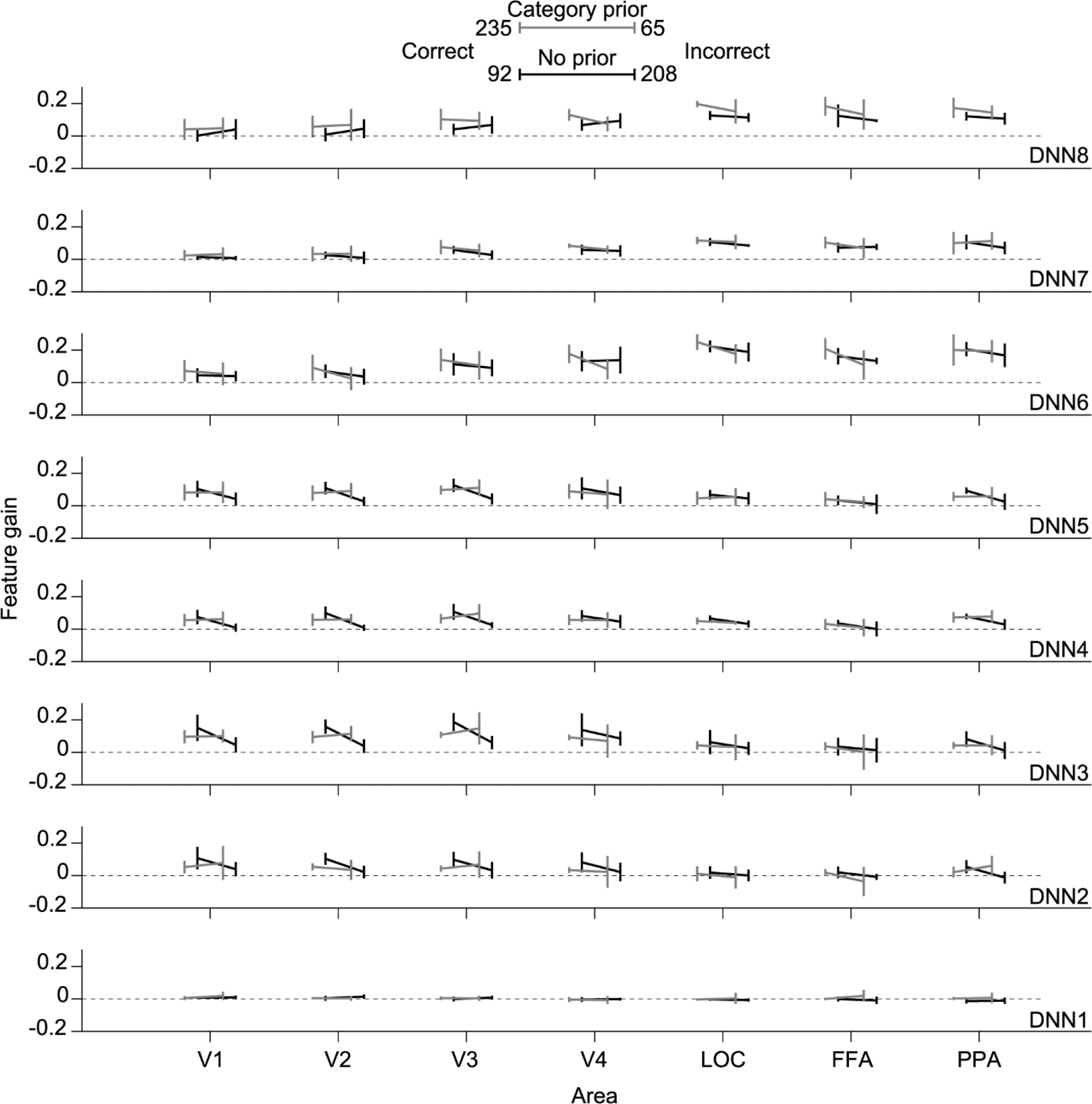
Effect of behavioral performance. Feature gain for features predicted from different visual areas grouped by experimental condition (category-prior vs. no-prior) and recognition (correct vs. incorrect). Legends include the total number of occurrences of each response across subjects. Mean feature gain is indicated for each DNN layer (blur levels pooled, 0% excluded). Error bars indicate 95% CI across five subjects.

We also analyzed our data by grouping it according to certainty level (certain and uncertain). We show the results of mean feature gain over subject means for each DNN layer of this analysis in Figure 7. For the category-prior condition, when an image was recognized with certainty we found significant enhancement in feature gain in DNN5, while for the no-prior condition significant enhancement was found in DNN1 and 7.

**Figure 7.**
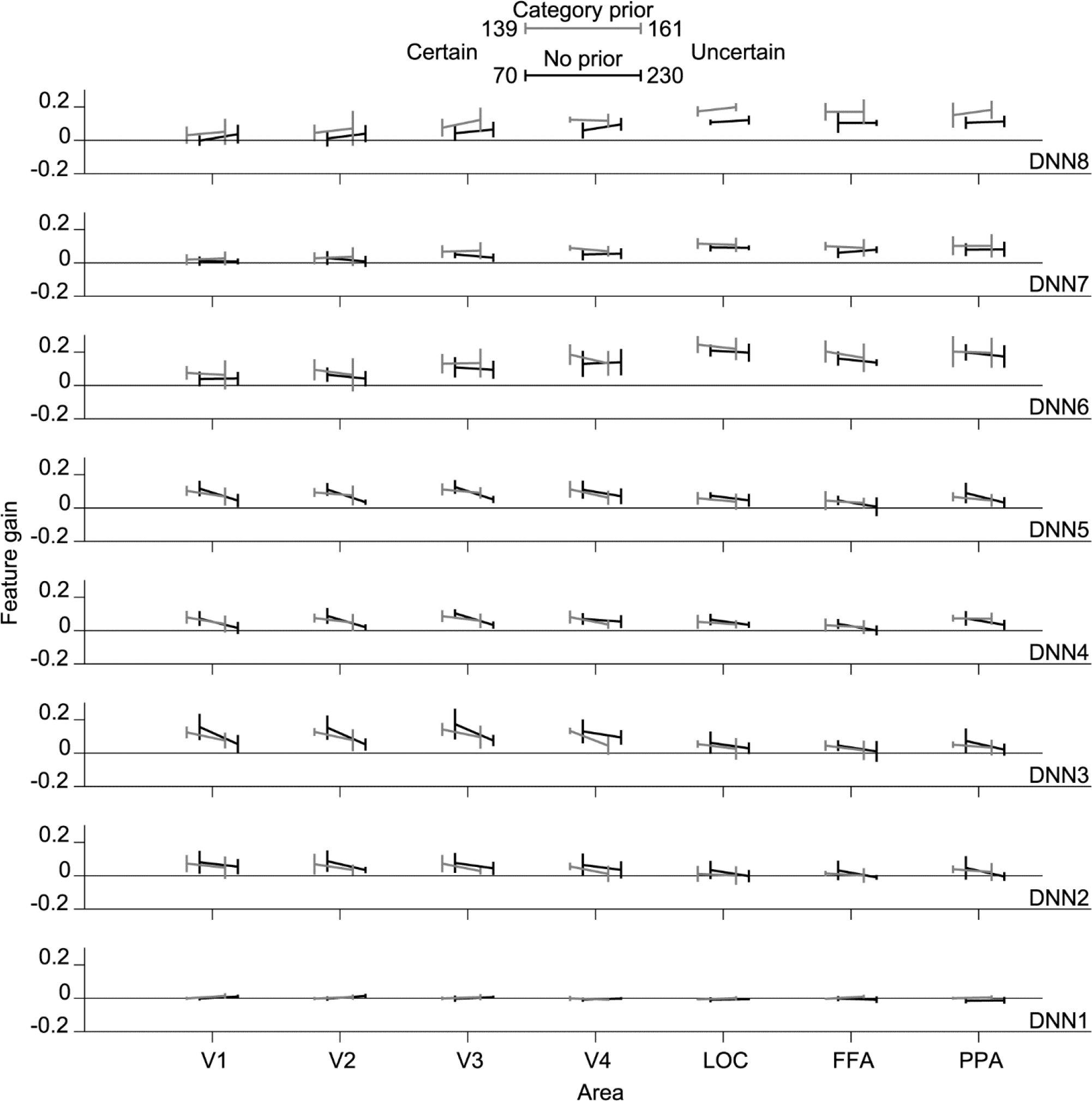
Effect of confidence level. Feature gain for features predicted from different visual areas grouped by experimental condition (category-prior vs. no-prior) and confidence level (certain vs. uncertain). Legends include the total number of occurrences of each response across subjects. Mean feature gain is indicated for each DNN layer (blur levels pooled, 0% excluded). Error bars indicate 95% CI across five subjects.

From the results of Figures 6 and 7, we can observe that in some layers and conditions, recognition has a significant boosting effect on feature gain. However, we also found a considerable feature gain even without recognition that indicates a sharpening effect not guided by subjective recognition. This could be caused by a lower-level sharpening associated with local similarity or object component sharpening that could be common across different objects (like body parts in animals).

## Discussion

In this study, we have demonstrated sharpening of the neural representations of blurred visual stimuli. This sharpening can assist the visual system in achieving successful prediction. It originates from endogenous processing elicited by top-down projections or recurrent connections (or both) in the visual cortex. Compared with pure-feedforward behavior, the neural representations of blurred images tended to be biased towards those of corresponding original images, even though the original images had not yet been viewed. This sharpening effect was also found to follow a visual hierarchy similar to that in the visual cortex. We found that this sharpening was content-specific, and not just due to a natural image bias. It was also shown to be boosted by giving category information to the subject prior to stimulus viewing. This indicated that adding a more specific prior leads to further sharpening of the neural representations. However, we did not find that recognition had a strong role in boosting the enhancement process.

In our experimental protocol, the subjects viewed blurred stimuli in randomly organized sequences. In each sequence, different levels of blur of the same image were shown, ordered from the most blurred to the non-blurred stimulus (Figure 1A). This ensured that subjects did not have pixel level information. Nonetheless, the results show a tendency for the blurred images’ neural representations to correlate with the original images (Figure 2A and B). Conversely, the feedforward behavior demonstrated by the noisy DNN output showed an opposite tendency (Figure 2C). We computed the feature gain to investigate how the predicted DNN features deviated from pure feedforward behavior. Feature gain analysis showed that the predicted features are rather correlated with the original image features (Figure 2E). This indicates that a sharpening effect happens across the visual cortex, leading to a more natural-image-like neural representation.

We notice also that feature gain is relatively higher in deeper layers of the DNN (DNN6– 8, Figure 2E). This effect could be caused by the nature of image degradation. Image blurring tends to conceal localized details in favor of the global shape information. This could lead to the subject attempting to recognize the global object while ignoring localized details. Another observation is that feature gain in deeper layers drops between the 12% and 6% blur levels. At the 6% blur level, localized details start to unravel in most of the stimulus images. This could cause the lower layers’ feature gain to increase at the expense of the deeper layers’ feature gain (Figure 2E). If we consider the stimulus sequence from the most blurred stimulus to the original image stimulus, we could visualize the time scale of the top-down effect where deeper layers peak earlier than shallower ones. This could be one effect of our image presentation protocol where the subject is accumulating evidence at each level starting from global shape evidence followed by localized details to confirm their concordance with the global shape evidence.

This representation was also confirmed as not being due to a natural-image bias caused by the decoder training dataset, which consisted of natural unaltered images (Figure 3). These results are in line with previous studies showing that neural representations are improved due to a top-down effect (Lee and Mumford, 2003; Hsieh et al., 2010; Kok et al., 2012; Gayet et al., 2017). Kok et al. (2012) demonstrated that even though the overall neural activation weakens, the neural representations improve when a stimulus is in agreement with the expectation. Gayet et al. (2017) also showed that visual working memory enhances the neural representations of viewed stimuli. The reverse hierarchy theory also suggests that top-down modulation serves to fine-tune sensory signals by means of predictions initially made using lower spatial frequency features (Hochstein and Ahissar, 2002; Ahissar and Hochstein, 2004). Furthermore, Revina et al. (2017) showed that blurred stimuli can generate top-down processes that generalize to higher spatial frequencies. Modelling studies that incorporated top-down and recurrent connections have also shown a sharpening-like effect under an image degradation scheme visible in text-based CAPTCHAs (George et al., 2017).

Our analysis also shows that feature gain follows a similar hierarchy to the visual cortex (Figure 4). This indicates that the sharpening process occurs in the same hierarchical processing localization as normal processing where low level sharpening occurs in lower visual areas and enhancement of higher level features occurs in the higher visual areas. It could be indicative of a convergent mechanism by which bottom-up and top-down pathways are integrated into a single neural representation of the stimulus. This suggestion could be supported by previous reports on the prediction of visual features. Horikawa and Kamitani (2017a) demonstrated that visual perception and mental imagery yielded feature prediction that was homologous with that of the visual cortex hierarchy. Horikawa and Kamitani (2017b) also showed similar results from dream induced brain activity. Earlier studies showed strong representational similarities between the deeper layers of DNN and the brain activity in the inferior temporal cortex (IT; Cadieu et al., 2014; Khaligh-Razavi and Kriegeskorte, 2014; Yamins et al., 2014; Güçlü and van Gerven, 2015). This could be further investigated by high resolution imaging to reveal the layered structures in the visual cortex and analyze the neural representations in each layer. We demonstrate that top-down effects also show similar homology, thus suggesting that DNN-based methods are useful for studying visual top-down pathways since it can reveal the localization of the sharpening by means of DNN layer feature gain.

When we added a category-prior to the task, the number of competing categories for recognition decreased, thus the subjects tended to have a more directed top-down effect, due to the fewer number of competing stimuli (Bar and Aminoff, 2003). This led to a higher feature gain that was especially noted in the higher layers (Figure 5). This further supports the idea of neural representation sharpening when given a prior describing the stimulus content, as the top-down signal would be more correlated with the correct recognition results, thus leading to a stronger feature gain.

We also found that when subjects successfully recognized the image content, the feature gain in some layers predicted from lower visual areas was significantly improved. However, this was not salient as a general trend (Figures 6, 7). It was expected that recognition would lead to a boost in feature gain from the sensory competition perspective, as the subject would attend to the successfully recognized object in the stimulus image, leading to a directed top-down effect (Moran and Desimone, 1985; Kastner et al., 1998, 1999). Hsieh et al. (2010) also showed that successful recognition of binary images leads to a neural representation that is more correlated with the natural image, although in their study, recognition was driven by ground truth image viewing before watching the degraded stimulus at a later time.

Our results could be justified by the findings in de-Wit et al. (2012), where top-down prediction was shown to be topologically inaccurate as it led to activity reduction in the whole of V1, rather than at the predicted location. Revina et al. (2017) also found that brain patterns due to top-down modulation did not share information with the corresponding bottom-up signals. In the prediction error realm, successful prediction would lead to zero error in the higher visual areas, and thus feature gain would decrease. Our results show an opposite, albeit weak effect, which nonetheless supports the representation-sharpening rather than prediction error hypothesis. Thus, prediction error mechanisms do not appear to be in operation when stimuli are blurred or they could be calculated and used as the sharpening signal as proposed in Kok et al. (2012). The sharpening effect without recognition may be driven by more localized and lower-level feature mechanisms. These mechanisms would enhance features corresponding to local components of the main objects that were common across many objects (i.e. eyes in animals). These local enhancement effects could lead to different recognition results. This was shown to be true for computer vision DNN-based deblurring algorithms, where the results of the enhancement process can lead to different results, according to the desired object (Bansal et al., 2017).

From these results, we can deduce that top-down modulation is in operation when visual input is degraded, even in the absence of a memory or expectation prior. Previous studies have proposed that the brain makes an initial processing step using low spatial frequency information. This step generates predictions of the content of the image in the orbitofrontal cortex; these predictions are then used to drive the top-down modulation effect (Bar and Aminoff, 2003; Bar et al., 2006; Kveraga et al., 2007; Breitmeyer, 2014). This top-down effect comes about in the form of sharpening of neural representations resulting from viewing degraded images. The mechanisms by which this effect materializes have been mostly overlooked in previous literature, due to the difficulty in finding a baseline for measurement. There has been more focus on the source of this top-down modulation effect than on how it materializes in the visual cortex (Bar et al., 2006; Chiou and Lambon Ralph, 2016). As we demonstrate here, the DNN representations could offer a plausible proxy for representing brain activity and for attaining a pure-feedforward baseline that can be used for measuring top-down effects. The illustrated enhancement was shown to be affected by the presence of prior semantic information, leading to a boost in the enhancement effect that was more visible in higher-level features. To the contrary, successful recognition did not also cause an overall boost in neural representation enhancement. Our results contribute to the long-standing question of how top-down and recurrent pathways affect bottom-up signals to achieve successful perception, which is believed to cause the hallucinatory symptoms associated with psychological disorders such as schizophrenia when their balance is disrupted (for review, see Friston et al., 2016; Jardri et al., 2016). Moreover, our stimulus presentation protocol could be used to test more comprehensive models of decision making under accumulation of evidence tasks (Platt and Glimcher, 1999). We have examined the question from a more general perspective of vision, which has allowed us to achieve a more comprehensive understanding of the vision process.

## Author contributions

MA and YK designed the study. MA performed experiments and analyses. MA and YK wrote the paper.

## Acknowledgements

The authors thank Tomoyasu Horikawa, Kei Majima, Mitsuaki Tsukamoto, and Shuntaro Aoki for help with data collection and analysis. We also thank the members of Kamitani Lab for their valuable comments on this manuscript. This research was supported by grants from JSPS KAKENHI Grant numbers JP15H05920, JP15H05710, and ImPACT Program of Council for Science, Technology and Innovation (Cabinet Office, Government of Japan), and the New Energy and Industrial Technology Development Organization (NEDO), and Japanese Government Scholarship (MEXT).

We thank Karl Embleton, PhD, from Edanz Group (www.edanzediting.com/ac) for editing a draft of this manuscript.

This study was conducted using the MRI scanner and related facilities of Kokoro Research Center, Kyoto University

## Conflict of Interest

The authors declare no competing financial interests.

## Funding sources

JSPS KAKENHI; JP15H05920; Mohamed Abdelhack, Yukiyasu Kamitani

JSPS KAKENHI; JP15H05710; Mohamed Abdelhack, Yukiyasu Kamitani

ImPACT Program of Council for Science, Technology and Innovation; N/A; Yukiyasu Kamitani

The New Energy and Industrial Technology Development Organization (NEDO); N/A; Yukiyasu Kamitani

Japanese Ministry of Education, Culture, Sports, Science and Technology Scholarship (MEXT); N/A; Mohamed Abdelhack

